# Sleep-wake Characteristics in a Mouse Model of Severe Traumatic Brain Injury: Relation to Post-Traumatic Epilepsy

**DOI:** 10.1101/2020.06.16.137034

**Authors:** Sai Sruthi Konduru, Eli P Wallace, Jesse A Pfammatter, Paulo V Rodrigues, Mathew V Jones, Rama K Maganti

## Abstract

**Study objectives:** Traumatic brain injury (TBI) results in sequelae that include post-traumatic epilepsy (PTE) and sleep-wake disturbances. Here we sought to determine whether sleep characteristics could predict development of PTE in a model of severe TBI.

**Methods:** Following controlled cortical impact (CCI), sham injury (craniotomy only) or no craniotomy (NC), CD-1 mice were implanted with epidural electroencephalography (EEG) and nuchal electromyography (EMG) electrodes. Acute (1^st^ week) and chronic (months 1, 2 and 3 after injury) 1-week long video-EEG/EMG recordings were examined for epileptiform activity. We analyzed sleep-wake patterns manually and extracted high amplitude interictal events from EEG using an automated method. Sleep spindles and EEG delta power were derived from non-rapid eye movement (NREM) sleep epochs. Brain CTs (computerized tomography) were performed to quantify the extent of brain lesions in cohorts of sham and CCI.

**Results:** Posttraumatic seizures were seen with CCI, whereas interictal epileptiform activity as well as sleep-wake disruptions (shorter wake or NREM bout lengths, shorter duration or lower power for spindles, and increased NREM EEG delta power) were seen in CCI and sham groups. No sleep feature predicted PTE. Follow up brain CTs showed a small lesion in the sham injury group suggesting a milder form of TBI that may account for their interictal activity and sleep changes.

**Conclusions:** In our model, interictal epileptiform activity and sleep disruptions resulted from CCI and sham and thus, sham injury was not an optimal negative control. Further work is necessary to determine the relationship between sleep-wake disturbances and PTE.

**Statement of significance:** Traumatic brain injury (TBI) results in sequelae such as post-traumatic seizures and sleep-wake disturbances but it is difficult to predict which individuals will develop these symptoms. Our study is novel in that we characterized epileptiform activity and multiple sleep characteristics in a mouse model of severe TBI (Controlled cortical impact-CCI) and explored whether any specific sleep disturbance can predict post-traumatic epilepsy. Specifically, post-traumatic seizures were seen after CCI only whereas epileptiform activity other than seizures as well as sleep-wake disruptions in mice that received a TBI and their sham injury controls. CT imaging showed that the sham injury group also had small brain lesions suggesting that a more optimal control in TBI research is to perform no craniotomy. No single sleep characteristic was predictive of post-traumatic epilepsy although NREM delta power was different in chronic recordings between TBI mice that developed seizures and those that did not. These studies are relevant to further research in TBI models, to develop a sleep biomarker for PTE. The work is also relevant to humans with TBI as monitoring sleep phenotypes may predict risk, but may also help develop therapies to prevent post-traumatic epilepsy.

## Introduction

According to CDC, there are over 5 million people with a traumatic brain injury (TBI)-related disability in the United States^1^. The World Health Organization reports that TBI results in major disability and is referred to it as a “silent epidemic” ^2^. TBI can lead to a wide range of sequelae that include post-traumatic epilepsy (PTE), sleep disorders, cognitive and motor deficits, and neurobehavioral complications such as mood disorders or post-traumatic stress disorder (PTSD)^3^. Of these sequelae, epidemiological data show that PTE is seen in about 10-20% of patients with TBI in the civilian population but as high as 50% of injured military personnel^4, 5^. The 30-year cumulative incidence of epilepsy is 2.1% for mild, 4.2% for moderate, and 16.7% for severe TBI^6^. In humans and animal models alike, post-traumatic seizures that develop in the first week are defined as acute post-traumatic seizures and those that develop later are defined as late post-traumatic seizures or PTE^7, 8^. In humans, latency to PTE can be months to years after injury ^7, 8^. Post-traumatic seizures can be unifocal (such as in the hippocampus or at the site of cortical injury), bilateral and rarely, generalized^9^. While some risk factors for development of PTE such as bilateral contusions, intracranial hemorrhage, penetrating injuries, older age, and prolonged coma after injury are known, it is currently very difficult to predict who will develop PTE after TBI.

Studies have shown that sleep-wake disturbances are quite common after TBI, but the clear link between TBI severity and the development of sleep-wake disturbances is not well established^10^. Surprisingly, sleep-wake disturbances may be more common after mild or repetitive mild TBI^11^. Sleep-wake disturbances can begin immediately after TBI, and persist long after sustaining the injury^12–13^. In humans, there is a spectrum of sleep complaints among TBI patients ranging from insomnia to hypersomnia^10^ with polysomnographic studies showing sleep fragmentation, increased awakenings after sleep onset, reduced REM sleep^13^, reduced non-rapid eye movement (NREM) sleep delta power and reduced latencies on multiple sleep latency tests ^14^. Similarly, in animal models of severe TBI, sleep disruptions are evident that include increased NREM sleep or shorter sleep/wake bouts and reduced time in REM sleep acutely^15^ or chronically^16^. Sleep-wake disturbances and PTE can occur in the same person and whereas several mechanisms of PTE have been elucidated^17^, it is not known whether they have shared mechanisms although there is evidence of both hippocampal and thalamic GABA_A_ receptor subunit changes^18^ that favor deficits in synaptic inhibition. Interestingly, sleep and epilepsy appear to have a synergistic relationship whereby epilepsy disrupts sleep and sleep disruptions are a common trigger for seizures.

Here, we hypothesized that TBI results in sleep-wake disturbances early on and that one or more sleep characteristics may serve as a biomarker for PTE. In a model of severe TBI (Controlled Cortical Impact, CCI), we studied epileptiform activity including electroclinical seizures, sleep architecture; characteristics of sleep spindles and NREM sleep delta power in acute (week 1) and chronic (months 1, 2 or 3) recordings. Our data show that acute or chronic post-traumatic seizures were seen in the CCI group only. However, visually identified interictal epileptiform activity, high amplitude interictal events identified from automated analysis, and sleep wake disturbances were seen both in CCI and sham injury groups. Sleep-wake disruptions consisted of shorter wake or NREM sleep bout lengths, lower amplitude or power for sleep spindles, and increased NREM delta power (a marker of sleep homeostatic pressure), in both sham and CCI groups compared to NC. No single sleep characteristic distinguished the CCI group or was predictive of PTE. A follow up imaging study then showed a well-defined lesion in CCI group but a small volume lesion in the area of craniotomy was seen in the sham injury group suggesting that they possibly had a milder form of brain injury.

## Methods

### Animals

Adult male and female CD-1 mice (~4 months old) were obtained from Jackson Laboratory (Bar Harbor, ME). Mice were group housed until surgery and were singly house during EEG recording days. Mice were housed under a 12:12 light:dark cycle at 29±1 °C with food and water provided ad libitum. All procedures involving the use of animals were approved by the University of Wisconsin IACUC in accordance with the US Department of Agriculture Animal Welfare Act and the National Institutes of Health policy on Humane Care and Use of Laboratory Animals.

### Induction of TBI and electrode implantation

Briefly, CCI was delivered (n=40) to induce a moderate-to-severe TBI to the right temporo-parietal cortex and underlying dorsal hippocampus. Under isoflurane anesthesia and in a stereotaxic frame, a 4-5 mm craniotomy was performed, centered at a location 5 mm posterior from bregma and 3 mm lateral. The dura was left intact. Isoflurane anesthesia was maintained at 2% for several minutes prior to delivery of the impact. Utilizing a Precision Cortical Impactor (Hatteras instruments, NC) with a 3 mm diameter tip, a moderate-to-severe impact (2 mm depth, 5 m/s impact velocity and 100-200 msec dwell time) was delivered. Following the CCI, epidural screw electrodes were placed in the right frontal (1.5 mm anterior and lateral to bregma) and left parietal area (1.5mm posterior and lateral to bregma) as well as occipital reference. Additionally, two stainless steel wires were placed in the nuchal muscles to measure EMG for sleep scoring. Electrodes were connected to a head cap, and the electrode-head cap assembly was fixed to the skull with dental acrylic. Sham injury animals (n=24) received anesthesia for a similar duration with a craniotomy but no CCI delivered but EEG and EMG electrodes were affixed as in TBI mice. A third control group (n=6) consisted of CD-1 mice that simply received anesthesia and EEG/EMG electrodes without a craniotomy.

Animals were allowed to recover for 1-2 days post-injury, with appropriate post-operative care. Animals were then transferred to individual recording chambers. Sequential twenty-four-hour video-EEG recordings were acquired with an Xltek amplifier (Xltek, Madison, WI), sampled at 1024 Hz and filtered between 1 and 70 Hz. During the recording, animals were housed under diurnal conditions (12 hours light and 12 hours dark), in individual Plexiglas chambers (10 inches tall and 6 inches in diameter). Continuous one-week long recordings occurred during the first week after surgery (acute) and then again (chronic) at month 1, 2 and 3 for TBI or Sham groups and at week 1 and month 1 for NC controls. Animals were housed in groups between recordings. The acquired data were converted to EDF format for review and analysis.

### Analysis of seizures and interictal spikes

EEG data were analyzed for seizures correlated with video. Apart from seizures, epileptiform activity consisted of isolated spikes (defined as isolated spike and wave of <200 milliseconds duration), spike trains or runs (defined as a rhythmic or semi rhythmic series of spikes lasting seconds or minutes), absence-like spike-wave discharges (defined as high amplitude 5-7 Hz events lasting 2-4 seconds) and frank electrographic seizures (Figure 1 B, C and D). The electrographic seizures (Figure 1A) were correlated with the behavioral characteristics on video and classified on a Racine scale. We used an automated method ^21, 22^ for the detection of high amplitude interictal events from EEG records of the same files in which sleep data were analyzed. Briefly, we identified high amplitude events (as starting when signal crossed above five standard deviations from the mean and ending upon crossing back below one standard deviation). Events were considered as discrete spikes (as opposed to being part of a seizure) if their durations were briefer than 200 ms and they were separated from other events by more than 200 ms. We then projected these events into Principal Components space and clustered events into nine groups (Figure 2E) using the first three Principal Components and a Gaussian Mixture Model. Within each cluster, we calculated the ratio of events belonging to animals from the TBI treatment to those from animals belonging to the CCI and NC treatments. This ratio (ranging from 0 to 1) provides the probability that any random event selected from within a cluster belongs to an animal that received CCI. From this analysis and for each EEG record we calculated the number of high-amplitude events per hour and the number of high amplitude events per hour with a ratio greater than or equal to 0.90 of being related to CCI (referred to as “CCI-related” events) (Figure 2 C and D). The significance of these latter events that are preferentially associated with TBI is that they may provisionally be considered as PTE-related epileptiform events (or interictal spikes in those animals that experienced seizures).

**Figure 1:**
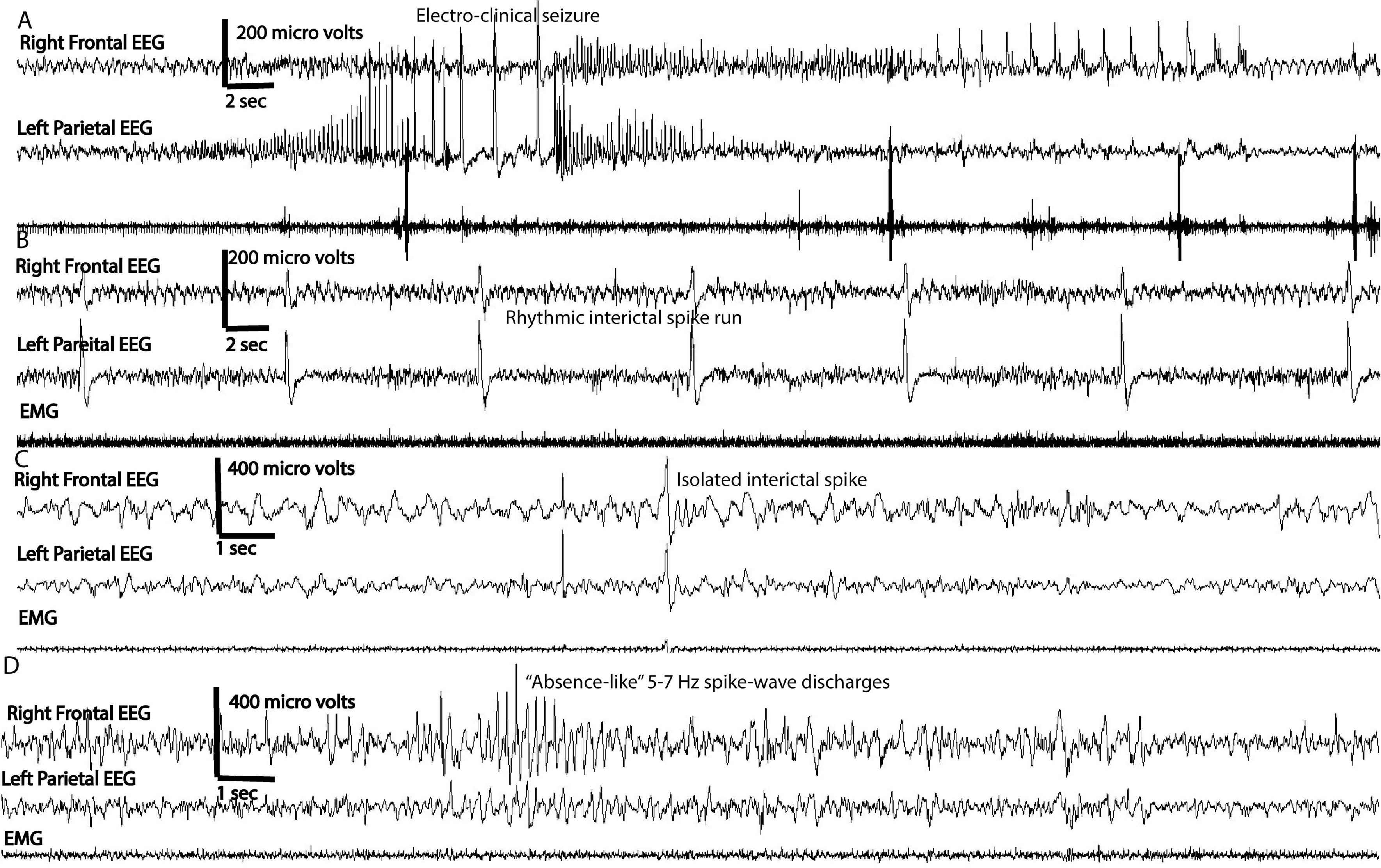
Epileptiform activity: Each animal had a right frontal and left parietal parietal EEG leads as well as an EMG electrodes. Panel A shows EEG tracing from an animal with an electro-clinical seizure where build up rhythmic spike-wave discharges is seen bilaterally. In Panel B a run of rhythmic or periodic interictal spikes is seen (60 second window) in a mouse with CCI. Panel C shows an isolated interictal spike in a mouse with CCI. Panel D shows an ”absence-like” spike-wave discharges that is characterized by high amplitude spikes occurring at ~5-6 hz lasting about 4 seconds.

**Figure 2:**
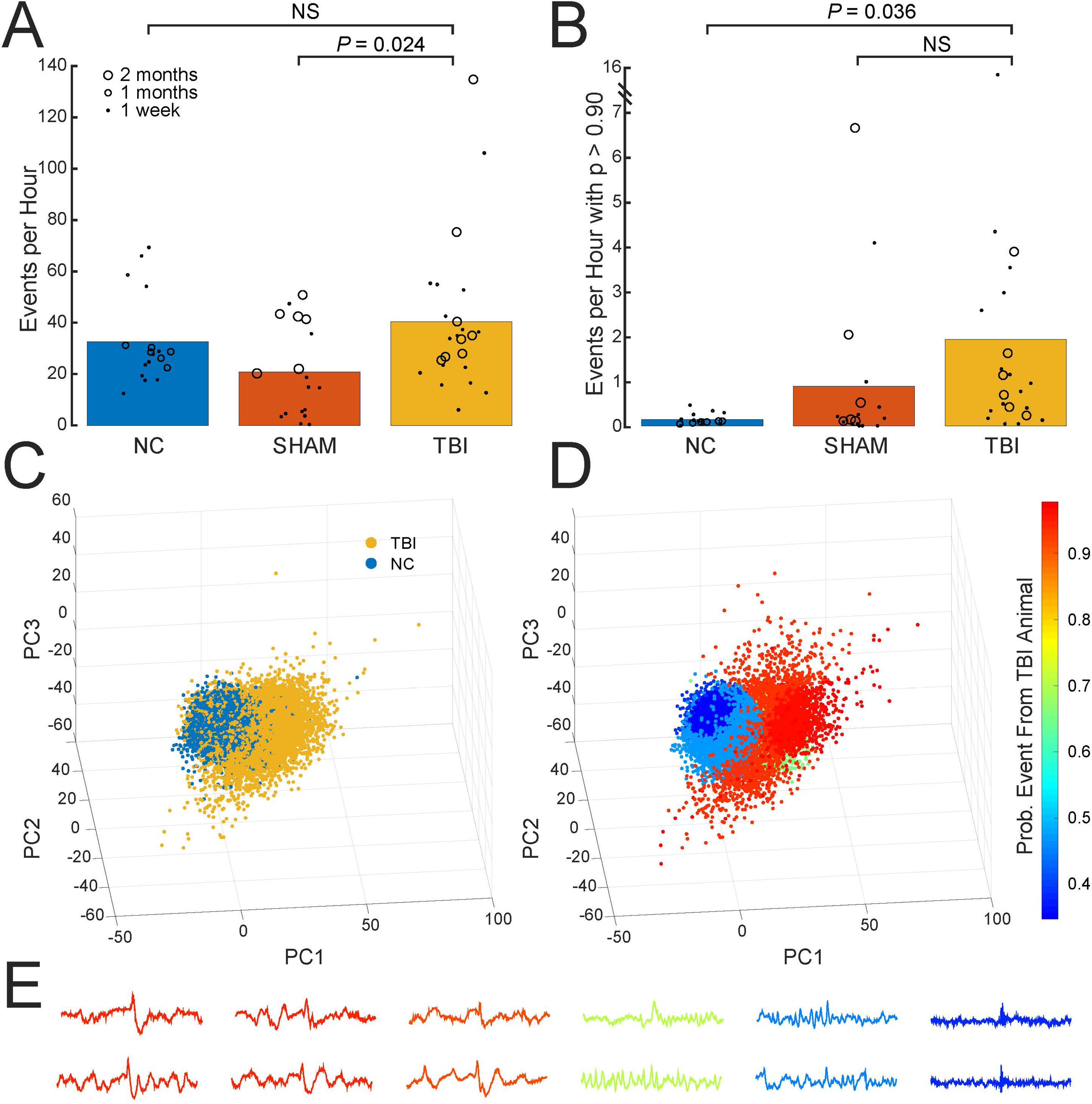
High amplitude event analysis: The frequency of high-amplitude events detected from NC, Sham, and TBI (CCI) EEG recordings (3A). There are certain types of events identified using the algorithm in Pfammatter et. al 2019 (those with a ratio greater than or equal to 0.90) that are predictive of TBI or TBI-like injury (Panel B). Panel C-D show how high-amplitude events were projected into Principal Components space and how the event clusters were formed how TBI-related ratios were applied to each cluster. Panel 3C shows the distribution of events from TBI and NC animals. The relative frequency of events from TBI animals to those from TBI and NC animals within each of nine clusters (PanelD) was used to identify events that have a high specificity (greater than 0.90, and those shown in Panel B) of being related to TBI treatment. Panel E shows a spectrum of events from those strongly related to TBI injury (left/red) to those less specifically related to the TBI treatment (right/green and blue).

### Analysis of vigilance states, sleep spindle characteristics and NREM delta power

EEG data were manually scored for vigilance states, in 4-second epochs using Sirenia Sleep (Pinnacle Technologies, Lawrence, KS), divided into Wake, NREM or REM according to EEG/EMG trace waveforms according to methods described previously^19^ (Figure S1). We scored sleep-wake patterns for days 4 and or 5 of each week-long recording (selected randomly), and all sleep analyses were done from these data. Manually scored sleep-wake patterns were available for 6 days in NC controls (from 3 mice), 7 days for Sham (from 7 mice) and 11 days in TBI group (from 9 mice) at week 1 and for 6 days (3 mice) in NC, 7 days (7 mice) for sham and 12 days (12 mice) in TBI group at month 1 or 2. Sleep parameters analyzed included total time spent in different vigilance states in 24-hours, vigilance state bout number (number of episodes of a given vigilance state across 24-hours) and vigilance state bout length. Sleep spindles were analyzed using automated algorithm with a code that was provided by Uygun et al 2019^20^, in MATLAB (Math Works, Natick, MA). The algorithm computes a root-mean-square (RMS) of the entire trace after band pass filtering at 9-15 Hz. The RMS values are then cubed to facilitate placement of thresholds. A selection threshold of 1.5 × cubed RMS and detection threshold of 3.5 times cubed RMS was used to detect spindles. An example of the algorithm output in a NC control is shown in Figure 3. The output from the original algorithm yielded mean spindle density (number of spindles per minute of NREM sleep); mean spindle duration (sec); mean spindle peak amplitude (μV), frequency (Hz) but was modified to yield spindle power (μV^2^). The algorithm was previously validated with comparisons to human scoring of EEG data though we have not performed independent validation^20^. In addition, EEG delta power was calculated for each scored NREM epoch by establishing power-spectral density with Fast-Fourier transformation and integrating the power spectral density from 0.5-4 Hz. To account for inter-animal variability, delta power values were normalized to the sum of power integrated over theta (5-9 Hz), sigma (10-14 Hz), and gamma (25-100 Hz) bands.

**Figure 3:**
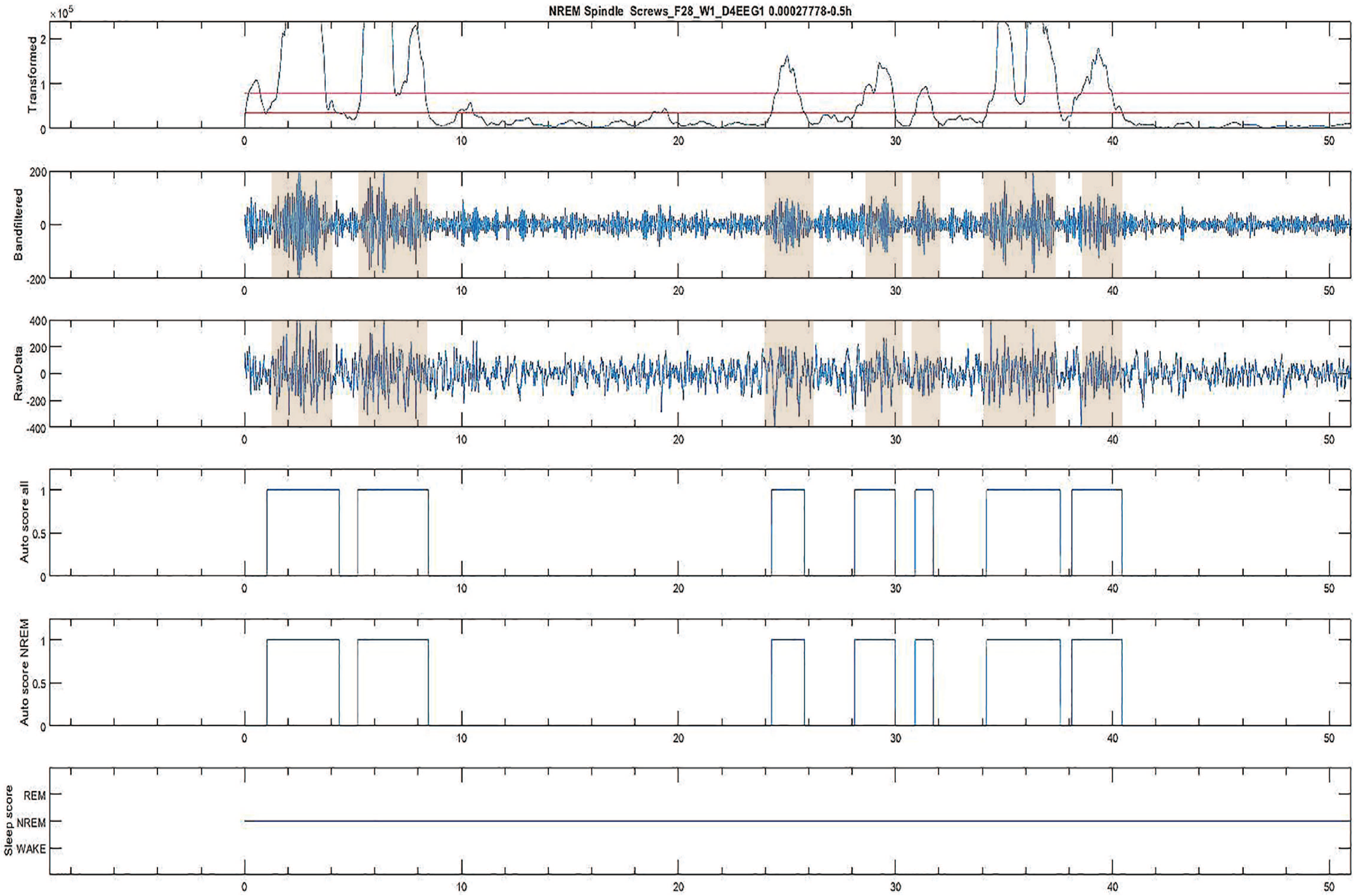
Spindle detection using a Matlab algorithm: This figure depicts sleep spindles output detected in one mouse through the automated MATLAB implemented algorithm. On the top row, cubed RMS signal with detection thresholds is seen. The 9-15 hz band pass filtered signal is seen in row 2 and the raw EEG signal is seen in row 3 band pass filtered between 1-30 Hz. A spindle is detected when it crosses the amplitude thresholds (highlighted in shaded area for both algorithm detected and a corresponding raw EEG detected spindle. Rows 4 and 5 show the duration of each detected spindle by the algorithm. Bottom row shows the stage of sleep. Shaded area shows the spindles on raw EEG as well as 9-15hz bandpass filtered EEG.

### Imaging

An imaging study was performed with computerized tomography (CT) scans in a subgroup of CCI and sham injury animals (n=5 each). Formalin-fixed mouse brains were CT scanned with a Siemens Inveon microCT (Siemens Medical Solutions USA, Inc., Knoxville, TN). All scans were acquired with the following parameters: 80 kVp, 1000 μA current, 700ms exposure time, 220 rotation steps with 600 projections, medium-high magnification, and binning factor of 2. Raw data were reconstructed with filtered back-projection applying the Shepp-Logan filter using the high-speed COBRA reconstruction software (Exxim Computing Corporation, Pleasanton, CA) yielding isotropic voxels of 31.5 microns. Analysis was conducted with Inveon Research Workplace (Siemens Medical Solutions USA, Inc., Knoxville, TN). We compared lesion size of any lesion identified in sham and CCI groups.

### Data Analysis

Time spent in different vigilance states as well as sleep efficiency across the 24 hours for NC, sham and CCI was assessed using ANOVA with post hoc Bonferroni-Holm corrections. Then, sleep-wake patterns or spindle density (defined as number of spindles per minute of NREM sleep) across the 24-hours was divided into different time bins (4-hour bins for sleep-wake and 6-hour bins for spindles) and within group or between group differences were evaluated using a mixed-model ANOVA. We used treatment (i.e. NC or Sham or TBI) as a fixed-effect variable and time as a random-effect variable. Data for means and 95% confidence intervals (CI) were extracted. Differences in amount of time spent in each vigilance state at each individual time bin was compared using one-way ANOVA with Bonferroni-Holm post hoc correction. Each group included in ANOVA analyses, data were not found to be skewed at a threshold less than [2]. NREM delta EEG power was analyzed after EEG data was subject to a Fast Fourier transformation (FFT) to extract discrete frequencies (in delta, theta, alpha, sigma and gamma bands) and binned into 4-hour blocks across the 24-hours. A mixed-model ANOVA analysis at each time point [acute (week 1) or chronic (month 1 or 2)] was performed, with Group (NC, SHAM, and CCI) as fixed-effect, and time bin (i.e., six 4-hour bins) as a random-effect. Multiple comparisons were completed with a Tukey-Kramer post-hoc correction. High-amplitude event counts were compared between TBI/NC and TBI/Sham treatment groups using two-sample t-tests with unequal variance. P-values were corrected by the Bonferroni method where each P-value was multiplied by the number of comparisons (two). Statistical analyses were conducted with an alpha value of 0.05.

## Results

### Seizures Following CCI

In acute or chronic video-EEG recordings, electro-clinical seizures were observed in about 23% (9 out of 40) of animals that had CCI but none were observed in the sham or NC controls. Among the CCI group 15 animals had both acute (Week 1) and chronic (months 1, 2 and 3) recordings and two of these (~13%) had acute post-traumatic seizures. Chronic only (months 1, 2 and 3) recordings were available for 25 CCI animals and of these, 7 (28%) developed PTE anywhere from month 1-3 after injury. One of the two that had acute post-traumatic seizures also had electro-clinical seizures during chronic recording. Only one animal out of 25 that had chronic recordings had seizures both at month 2 and 3. Thus, the total number of Racine grade 3 to 5 seizures was relatively small. The behavioral correlate for the electrographic seizure consisted of behavioral arrest, head nodding and hind limb stretching (Racine grade 2 to 3) that sometimes progressed to Racine class V seizures. The electrographic characteristics of the seizures consist of rhythmic fast activity that built into rhythmic spike wave discharges progressing bilaterally, similar to what was described previously by others (Figure 1A). Duration of the electro-clinical seizures varied from 20-160 seconds. No electro-clinical seizures were observed in sham or NC groups.

### Epileptiform Activity Other Than Seizures

Apart from the electro-clinical seizures, which were seen in a proportion of CCI animals, visual inspection of EEG traces showed epileptiform activity in a larger proportion of CCI animals (15 out of 15 in acute recordings and 17 out of 25 in chronic recordings). However, we observed these in the sham group as well. This activity consisted of semi rhythmic or rhythmic spike runs (Figure 1B) or isolated spikes (Figure 1C) or absence-like spike-wave discharges (Figure 1D). The spike runs and absence-like spike-wave discharges were not associated with any clear behavioral correlate such as freezing or wet dog shakes on video and thus, were not counted as electro-clinical seizures. The NC controls were not observed to have any form of epileptiform activity on visual inspection. We then performed an automated analysis of high amplitude interictal events with the notion that interictal epileptiform spiking appears as a high amplitude event compared to the rest of the background in EEG traces of humans or animals with an epileptic insult^21^. When comparing all the high-amplitude events, we found no difference between the NC and CCI groups (t(37.31)=0.525, uncorrected *p*=0.603, corrected *p*>1) but, CCI had a greater number of such events compared to sham (t(35.55)=2.63, uncorrected *p*=0.012, corrected *p*=0.024), (Figure 3A). However, when comparing high-amplitude events whose waveforms were most predictive of whether the animal received CCI (those with a ratio of greater than 0.90^21^), we found that the CCI group had a greater number of these events compared to NC (t(21.07)=2.58, uncorrected *p*=0.018, corrected *p*=0.036), but not compared to the sham group (t(34.48)=1.29, uncorrected *p*=0.205, corrected *p*=0.409) (Figure 3B). Therefore, compared to NC, both CCI and sham displayed a preponderance of events indicative of having received CCI.

### Vigilance State Characteristics

The 24-hour sleep efficiency (percent time spent sleeping during the 24-hour recording) was no different between the groups at week 1 (ANOVA: NC: 51.39±5.11; sham: 58.92±8.18; CCI: 52.60±8.07; F(2, 21) = 1.688; *p*=0.20). At month 1 or 2 however ANOVA with post-hoc Tukey test showed that sham group had greater sleep efficiency than CCI (NC: 55.59±2.80; sham:60.44±5.30; CCI: 51.48±2.33; F(2, 22) = 9.041; *p*=0.001). We then analyzed differences in the total time spent in different vigilance states (wake, NREM and REM) across the 24-hours (shown in minutes) in acute and chronic recordings. At week 1, there were no differences in time spent in wake or NREM between NC vs sham injury vs CCI groups (wake: F(2, 21)=1.78, *p*=0.19; NREM: F(2, 21)=0.7115, *p*=0.50). However, there was a difference in time spent in REM (F(2, 21) =9.16, *p*=0.001). Post-hoc Bonferroni-Holm correction showed that sham group had greater time spent in REM (121.18±28.42) compared to NC (75.28±18.60, p<0.05) or CCI groups (64.25±13.20, *p*<0.05) (Figure 4A). At month 1 however, there were differences in time spent in wake (F(2, 22)=10.19, *p*<0.001) as well as NREM (F(2, 22)=8.29, *p*=0.002) but not in REM (F(2, 22)=1.49, *p*=0.24). Post-hoc comparisons showed that Sham animals only had less time spent in wake (565.6±52.51) compared to NC (638.35±32.38, *p*<0.05) or TBI (677.76±20.66, *p*<0.01) and greater time spent in NREM (782.98±60.82) compared to TBI (677.72±25.11, *p*<0.01) (Figure 4B). Thus, no differences were seen in overall 24-hour sleep-wake patterns between NC controls and TBI at week 1 or month 1 or 2.

**Figure 4:**
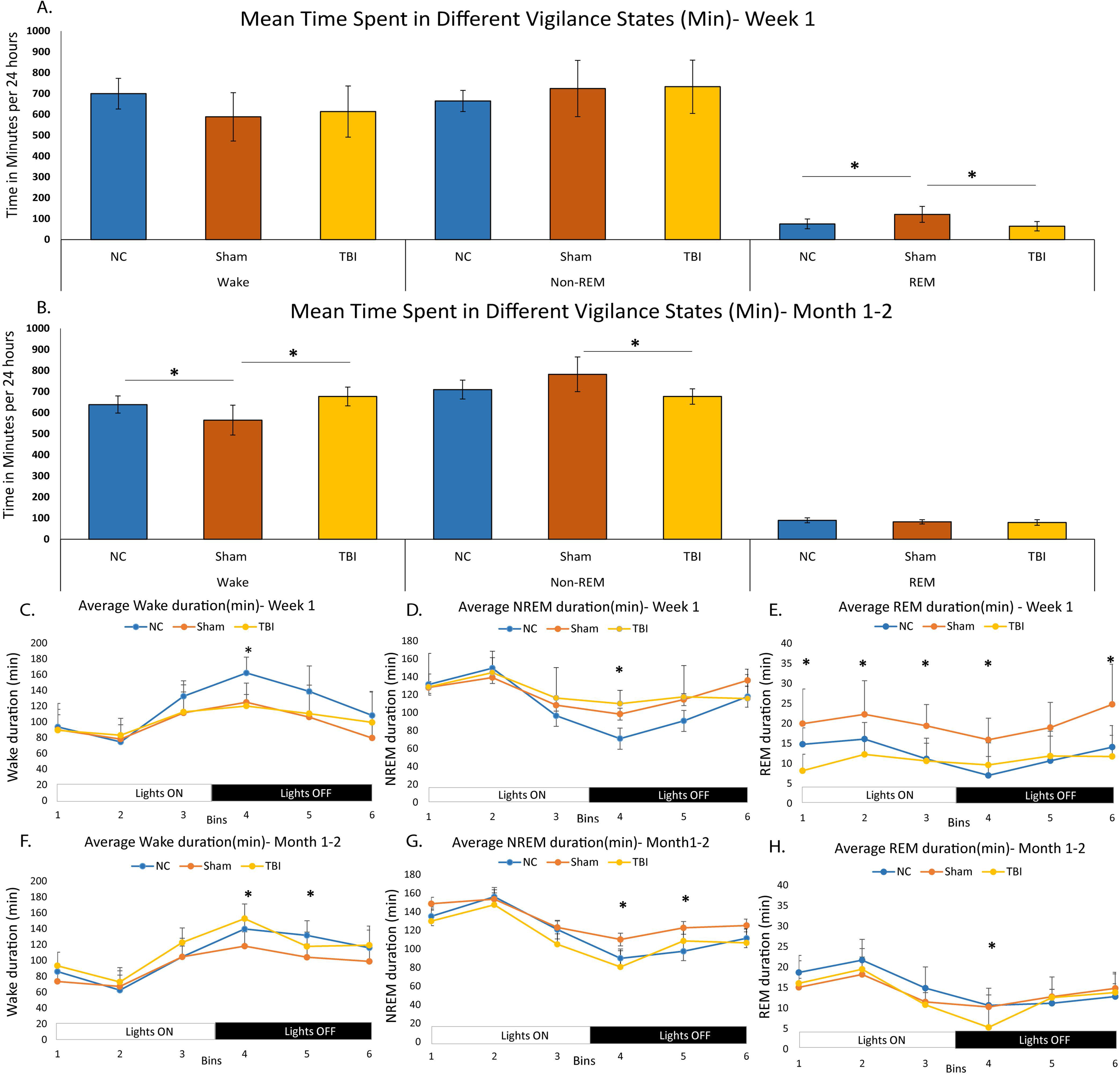
Vigilance state analysis: Figure 4 shows mean time spent in different vigilance states across 24 hours for Week 1 (A) and month 1 or 2 (B). Panels C to E shows oscillation of different vigilance states across 24 hours when examined in 4-hour bins. Differences in average times spent in Wake, NREM and REM in each bin is shown in minutes at Week 1 (Panels C, D and E) and at month 1 or 2 (Panels F, G and H). Significance seen in any measure is indicated by an *. Acutely, NC (blue) had more time in awake in the 4^th^ bin (6:30 pm to 10:30 pm), compared to Sham (red) or CCI (orange) (Wake: NC-162±8.23; Sham-124.74±9.17; TBI-120.02 ±4.38, F(2, 23)=10.03, *p*<0.001) and CCI had greater time in NREM (NC- 71.05± 6.55; Sham- 98.41 ± 10.42; CCI- 110.01±4.50, F(2, 23)=7.694, *p*=0.003). Panel E shows time spent in REM (in minutes) at week 1 between the 3 groups (Bin 1: NC- 14.78 ±1.69 Sham- 19.93±3.28 CCI- 8.17±1.24, F(2, 23)=9.073, *p*=0.001; Bin 2: NC- 16.09±1.69 Sham- 22.26±3.18 CCI- 12.26 ±1.11, F(2, 23)=6.93, *p*=0.004; Bin 3: NC- 11.13±1.60 Sham- 19.39±2.02 CCI-10.62 ±1.71, F(2, 23)=6.72, *p*=0.005; Bin 4: : NC- 7.00±1.92 Sham- 15.90±2.05 CCI- 9.60±1.66, F(2, 23)=5.04, *p*=0.01; Bin 6: NC- 14.08±2.20 Sham- 24.75±3.79 CCI- 11.73±1.59, F(2, 23)=7.75, *p*=0.003) (Figure 4, E). Chronically, Wake time was less for sham compared to CCI (in 4^th^ bin) or NC (in 5^th^ bin) (Bin 4- CCI: 152.68±5.35 vs NC: 139.53±4.44; Sham 117.90±6.88, F(2, 24)=9.25, *p*=0.001; Bin 5- Sham:122.61±6.15; NC: 97.45±6.59; CCI: 108.54±3.95, F(2, 24)=4.61, *p*=0.02). Sham had greater time spent in NREM sleep compared to CCI (4^th^ bin) or NC (in 5^th^ bin) (Bin 4- Sham: 110.12±6.99; NC: 89.86±3.08; TBI 80.57±3.95, F(2, 24)=7.50, *p*=0.003; Bin 5-Sham: 122.61±6.15; NC 97.45±6.59; TBI: 108.54±3.95, F(2, 24)=4.61, *p*=0.02). Time spent in REM was less for TBI compared to both NC or Sham in the 4^th^ bin (TBI: 5.30±1.33; NC: 10.68±1.70; Sham: 10.29±1.10, F(2, 23)=4.74, *p*=0.01).

When vigilance states were divided into six 4-hour bins across 24 hours (starting at 6:30 am or lights on), mixed model ANOVA showed a main effect of time where there was an oscillation seen in sleep-wake pattern with greater sleep during light-on and greater wake during lights-off (Figure 4 C, D, E) (data shown for wake in Table S1). We then compared differences in time spent in wake, NREM or REM among NC, sham and CCI groups in each of the 6 bins across 24 hours. The results seen did not show a consistent pattern of sleep-wake disruptions between the groups, but the only difference seen was in the 4^th^ time bin (6:30 pm to 10:30 pm which is the first 4 hours of “lights off”) in week 1 where NC group spent more time awake and CCI group spent more time in NREM. The latter is indicative of early morning sleepiness in CCI group (Figure 4 C and D). However, the time spent in REM was much greater in sham compared to NC or CCI in multiple time bins (Figure 4 C, D, E). At month 1 or 2, result was mixed with sham animals having more NREM sleep and less wake in the 4^th^ and 5^th^ bins which is 6:30 pm to 02:30am (Figure 4 F and G) and CCI has less time spent in REM compared to both NC or sham in the 4^th^ bin (Figure 4H).

Next, we evaluated the number of episodes of each vigilance state per 24 hours of recording termed “bout number” but no consistent pattern was seen. At week 1, sham groups had greater wake bout number than NC and greater NREM as well as REM bout number than NC or CCI (Figure S3 A-C). At month 1, the only difference seen was that CCI had greater number of wake bouts compared to either NC or sham (Figure S3 D-F). Lastly, we also compared the average “bout length” in seconds of wake, NREM or REM for each 24-hour recording at both week 1 and month 1 or 2 between the three groups. Analysis showed that unlike in the bout number, sham and CCI has shorter wake, NREM and REM bout length in both acute and shorter wake and NREM at chronic time points compared to NC (Figure S A-F). No difference was seen in bout lengths between sham and CCI groups.

When the TBI group was subdivided into those with or without post-traumatic seizures, none of the vigilance state characteristics described above stood out as a feature that is associated with presence of post-traumatic seizures.

### NREM Delta Power Analysis

Normalized EEG delta power was calculated via FFT in 4-second epochs of each 24-hour recording. Periods of NREM sleep, as determined by manual scoring of each epoch, were indexed and associated EEG delta power was parsed into six 4-hour bins and grouped by treatment and time of recording acute (week 1) or chronic (1 or 2 months post-surgery). Within the first week after surgery, we observed an increase in mean group NREM EEG delta power that followed relative severity of injury: NC had the lowest, sham had elevated NREM delta power over NC, and CCI animals had the greatest NREM delta power (Figure 5A, Table S3C, F(2, 10)=326.7, p<0.05). At the chronic time points, mean group NREM delta power of CCI animals decreased to levels similar to sham animals, while both sham and CCI animals exhibited greater NREM delta power than NC control animals (Figure 5B, Table S3C, F(2, 10) =168.49, *p*<0.05). We also assessed whether the development of post-traumatic seizures in CCI animals at any time during acute or chronic recordings differentially affected characteristics of NREM delta power. We observed post-traumatic seizures in 28% (4 of 14) of CCI animals included the analysis. At the acute time point, we observed no statistical difference between CCI animals based on development of seizures (Figure 5C, Table S3F, F(1, 5)=0.43, p>0.05). However, at the chronic time point, we found that CCI animals with detected seizures had lower NREM delta power than those CCI animals without detected seizures (Figure 5D, Table S3F, F(1, 5)=146.07, p<0.05). These findings suggest that acute increases in NREM delta power might be a useful diagnostic tool for TBI incidence or severity, as sham and CCI both exhibited elevated NREM delta power over NC controls and seemed to be indicative of TBI severity. Acute NREM delta power may be a poor predictor of whether animals with severe-TBI (CCI) would develop post-traumatic epilepsy, while prolonged monitoring of NREM delta power may provide insights into the likelihood of developing PTE.

**Figure 5:**
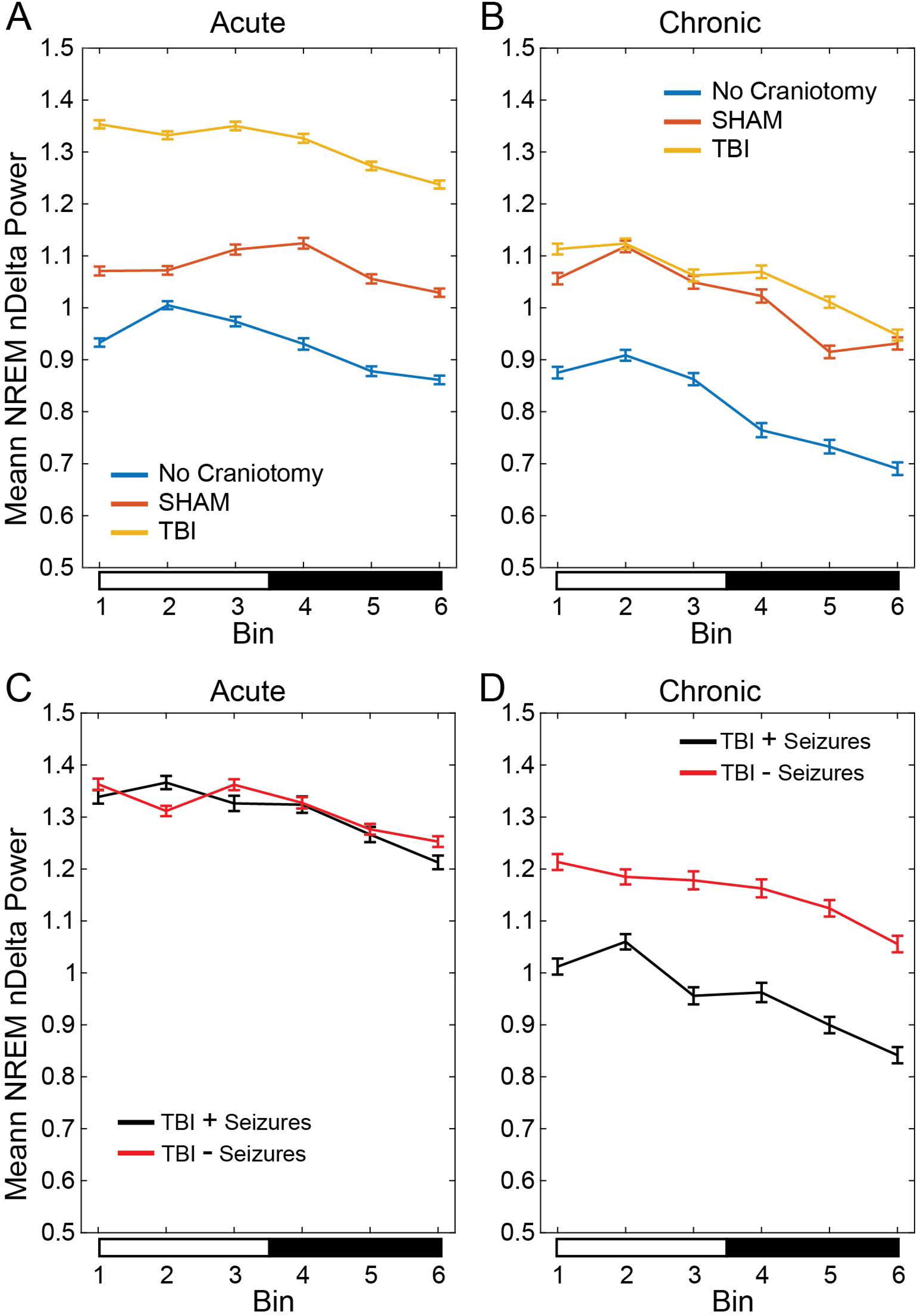
NREM delta power analysis: NREM delta power from EEG recordings collected from animals that had sham, or CCI, or no craniotomy controls across 24-hours. Each recording was parsed into 4-second epochs which were manually scored for vigilance state (wake, NREM, or REM) and transformed into power spectral density plots. EEG power were calculated for delta (0.5-4 Hz), theta (6- 9 Hz), sigma (10-15 Hz), and gamma (25-100 Hz) frequency bands. Delta power values were normalized to the sum power of delta+theta+sigma+gamma bands, and those normalized delta (nDelta) values associated with NREM sleep were isolated and further parsed into 6, 4-hour bins for each day of recording. Mixed-effect 2-way ANOVAs with Tukey-Kramer corrected post-hoc multiple comparison tests were run for recordings grouped by time relative to surgery, within 1 week of surgery (acute) or 1-2 months following surgery (chronic). Binned NREM nDelta power of no craniotomy (NC, EEG only, blue) controls, SHAM (craniotomy+EEG, red), and CCI (craniotomy+CCI+EEG, yellow) were tested at acute (panel A) and chronic (panel B) time points and graphed as mean ± 95% confidence interval (95CI). At the acute time point (panel A), analysis of NREM delta power showed a main effect based on time bin (F(5, 367606) =281.27, *p*<0.05, Table S3B) and surgical group (F(2, 10)=326.74, p<0.05, Table S3C), though no significant interaction was seen (F(10, 367606)=0.57, *p*>0.05). At the chronic time point (panel B), mixed-effects ANOVA revealed a main-effect of time (F(5, 201367) =571.31, *p*<0.05, Table S3B) and surgical group (F(2, 10)=168.49, *p*<0.05, Table S3C), though no significant interaction was seen (F(10, 201367)=0.87, *p*>0.05). To determine if the presence of post-traumatic seizures differentially affected NREM nDelta power characteristics, we split those in the TBI group that exhibited seizures (TBI + seizures, black) and those that did not (TBI – Seizures, red). Again, binned NREM nDelta power was tested with a mixed-effects 2-way ANOVA with Tukey-Kramer corrected multiple comparisons, grouped by presence or absence or seizures, at acute (panel C) and chronic (panel D) time points. At the acute time point (panel C), analysis of NREM delta power showed a significant effect of bin (F(5, 138115)=114.17, *p*<0.05, Table S2E), but not presence of seizure (F(1, 5)=0.43, p>0.05, Table S2F), or interaction (F(5, 138115)=0.68, *p*>0.05). At the chronic time point (panel D), mixed-effects ANOVA showed a significant effect of time bin (F(5, 75405)=153.07, p<0.05, Table S2E), seizure presence (F(1, 5) =146.07, *p*<0.05, Table S2F) and interaction (F(5, 75405) =13.26, *p*<0.05). Lighting condition is shown below each graph as lights-on (open bar), and lights-off (closed bar).

### Sleep Spindle Analysis

First, we evaluated spindle dynamics across the 24 hours by examining the average number of spindles per minute (spindle density) per each 6-hour bin of NREM sleep. Mixed model ANOVA showed a main effect of time. In addition, the spindles oscillate across the 24-hours, gradually increasing during lights on and gradually decreasing during lights off. However, there was no effect of treatment suggesting that no difference in spindle density was seen between the groups in any time bin (Figure 6A) (Table S2).

**Figure 6:**
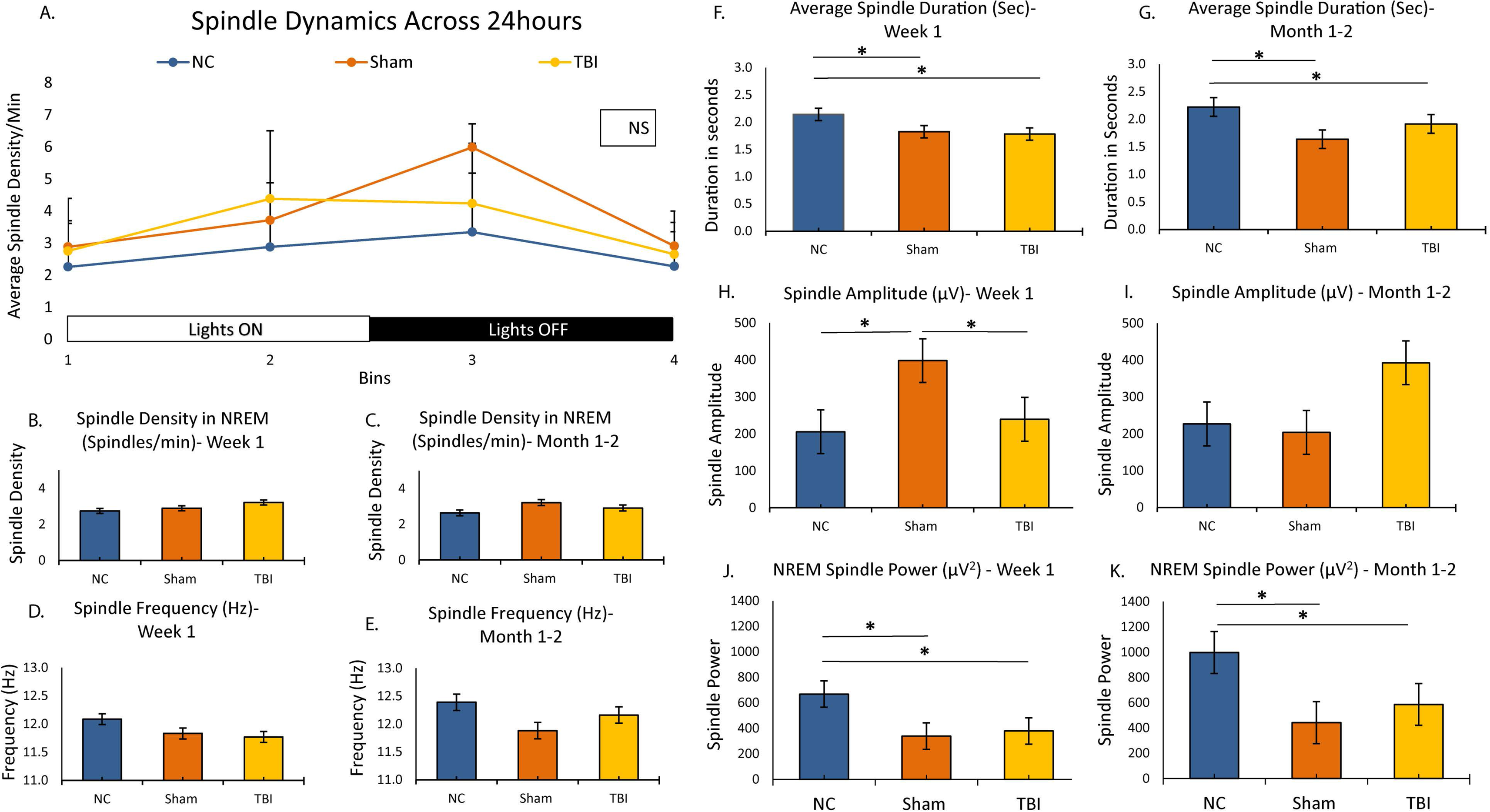
Sleep spindle analysis: Spindle characteristics are shown for NC, Sham and TBI animals at Week 1 or month 1 or 2. Spindle dynamics are shown in panel A where mixed model ANOVA showed a main effect of time (F (2 21)=8.02, *p*=0.001) but not of treatment group (F (2, 21) = 1.45; p=0.63). We also compared differences in spindle density between NC, Sham and TBI in each of the 4 bins using one-way ANOVA, no differences were found (Bin 1 *p*=0.60; Bin 2 *p*=0.18; Bin 3 *p*=0.13; Bin 4 *p*=0.63). Differences in other spindle characteristics at week 1 and month 1 or 2 are then shown for: spindle duration (panels B and C); spindle frequency (panels D and E); spindle amplitude (panels F and G), spindle density (panels H and I) and spindle power (panels J and K). Any significant difference between groups is indicated by an *. ANOVA analysis showed a main effect of group for spindle duration (in seconds) at week 1 (F(2 28)=5.58, *p*=0.009) and at month1 or 2 (F (2 17)=8.364, *p*=0.003); spindle power at week 1 (F (2 28)=10.959, *p*<0.001) and month 2 (F (2 17)=6.52, *p*<0.001) and spindle amplitude at week 1 only (F (2 28)=5.30, *p*=0.01). On post-hoc comparisons, at week 1, spindle duration in seconds was lower in sham (1.83±0.08, *p*<0.05) or TBI (1.78±0.09, *p*<0.05) compared to NC (2.14±0.08) but no difference was seen between sham and/or TBI. Similarly, at month 1 or 2, spindle duration was lower for sham (1.64±0.07, *p*<0.01) or TBI 1.92±0.08 (*p*<0.05) compared to NC (2.22±0.11) but no difference was seen between sham and TBI (*p*=0.15). For spindle power, post-hoc comparison showed that NC (668.99±69.05μV^2^) had higher power than Sham (339.21±49.63μV^2^, *p*<0.01) or TBI (379.26±33.38 μV^2^, *p*<0.01) at week 1 and at month 1 or 2 also (NC: 996.92±100.44μV^2^; Sham: 442.78±115.99 μV^2^, *p*<0.05; or TBI 586.52±88.33 μV^2^, *p*<0.05), but there was no difference between Sham and TBI either at week 1 or month 1 or 2.

We then examined other NREM spindle characteristics including spindle duration, spindle frequency, amplitude and power (data extracted for first 12 hours or lights on) both at week 1 and at month 1 or 2 of recordings among the 3 groups. Among these, spindle duration and spindle power were lower in CCI and sham compared to NC in both acute or chronic recordings (Figure 6 F and G; J and K respectively). Spindle amplitude was also lower in CCI and sham compared to NC at week 1 (Figure 6 H) but not at month 1or 2 (Figure 6 I). No differences were in other spindle characteristics between the groups (Figure 6 B-E and L) (Table 1). When CCI group was divided into those with and without post-traumatic seizures, none of the spindle characteristics stood out as a distinguishing feature in animals that developed post-traumatic seizures acutely or chronically.

**Table 1:** Analysis of different spindle characteristics: Table shows the different spindle characteristics we analyzed including density duration, frequency, amplitude and power. Differences between the groups was analyzed with ANOVA and post-doc Bonferroni-Holm correction. The p values from ANOVA were shown the table. ON post-hoc corrections, spindle duration was higher in NC compared to sham (p<0.01), and CCI (p<0.05) but not different between sham and CCI (p=0.89) at week 1 and month 1 or 2. Similarly, spindle power was higher in NC control compared to sham (p<0.01) or CCI (p<0.01) but CCI was not different from sham (p=0.63) at week 1 or month 1 or 2. Spindle amplitude was lower in NC compared to sham (p<0.05) or CCI (p<0.05) at week 1 only. No differences were seen between the groups in spindle density or frequency at week 1 or month 1 or 2.

| Table 1: Spindle characteristics in different groups, analyzed with ANOVA |  |  |  |  |  |
| --- | --- | --- | --- | --- | --- |
| Week 1 | Density<br>(number per<br>min)<br>Mean±SD | Duration (in<br>seconds)<br>Mean±SD | Frequency<br>(in Hz)<br>Mean±SD | Amplitude (in<br>μV) Mean±SD | Power (μV<sup>2</sup><br>Mean±SD) |
| NC Control | 2.74±0.48 | 2.14±0.28 | 12.09±0.30 | 205.33±34.74 | 668.99±239.22 |
| Sham Injury | 2.89±0.70 | 1.83±0.21 | 11.83±0.37 | 397.62±215.09 | 339.21±131.31 |
| TBI | 3.21±0.56 | 1.78±0.31 | 11.77±0.39 | 397.62±215.09 | 379.26±115.65 |
| F(2, 30) | 2.104 | 5.58 | 2.53 | 5.30 | 10.95 |
| P value | 0.14 | 0.009 | 0.09 | <0.001 | <0.001 |
| Month 1 or 2 |  |  |  |  |  |
| NC Control | 2.63±0.62 | 2.22±0.27 | 12.39±0.39 | 226.41±36.36 | 996.92±246.04 |
| Sham | 3.21±0.35 | 1.64±0.17 | 11.88±0.51 | 203.47±62.22 | 442.78±259.37 |
| TBI | 2.91±0.37 | 1.92±0.23 | 12.16±0.39 | 392.48±212.64 | 586.52±265.01 |
| F(2, 19) | 2.141 | 8.36 | 1.90 | 3.44 | 6.52 |
| P value | 0.14 | 0.003 | 0.17 | 0.055 | 0.007 |

In summary, analysis showed no consistent sleep-wake patterns distinguished CCI from other groups. The main observations that are consistent in acute or chronic recordings were that both sham and CCI had shorter wake and NREM bout length, shorter spindle duration and lower power and increased NREM delta power compared to NC control.

### Imaging data

The interictal epileptiform activity as well as sleep disruptions seen in both CCI and sham groups, led us to not only perform imaging study, but also employ NC controls during this study. Analysis of the CT imaging in CCI and sham injury groups (n= 5 each) showed a well-defined lesion in CCI group, but the sham group also had a lesion that is visually evident to the extent of the craniotomy on the images (Figure S4). However, the lesion volume was clearly larger for the CCI group compared to sham (CCI: ± 7.2 mm^3^, sham: 0.82 ± 0.68mm3; *p* = 0.039) (Figure S4) seen on the side of injury or craniotomy. What is evident from imaging is that the sham group likely had a milder form of brain injury that could have occurred during the craniotomy.

### Correlations of High Amplitude Events to Sleep-wake Patterns

We then examined whether interictal events contributed to sleep disruptions by correlating interictal high amplitude events per hour of recording with time spent in Wake in each hour in the three groups. We found no correlation (Pearson correlations: NC: r^2^ = 0.25, p= 0.31; sham: r^2^ = 0.07, p = 0.79; CCI: r ^2^= 0.13, p = 0.60), suggesting that interictal high amplitude events themselves did not correlate with total Wake time.

## Discussion

Overall, the key findings in our model of CCI are: a) following CCI, acute post-traumatic seizures were seen in 13% and late posttraumatic seizures (PTE) in 28% of animals; b) whereas no seizures were seen in sham and NC controls, epileptiform activity, high amplitude interictal events and sleep-wake disturbances were seen in both sham and CCI groups; c) sleep-wake disturbances in acute and chronic recordings consisted of shorter wake and NREM sleep bout lengths, shorter duration or lower power for sleep spindles and increased NREM delta power in CCI and sham compared to NC; d) no sleep characteristic predicted PTE; and e) finally, CT imaging showed that CCI animals had a large volume brain lesion whereas sham injury animals had a small but definite lesion at their craniotomy site suggesting that they had at least a mild form of brain injury.

Our data are consistent with several prior studies that demonstrated the presence of spontaneous seizures in the CCI model. Seizures were reported anywhere from 9% to as much as 50% of animals following CCI^23–26^. Seizures were reported as early as within 24-hours of impact^25–27^ or late as 6-9 months after injury^23^. Latency to spontaneous seizures was weeks to months in all the studies^17^. We found acute post-traumatic seizures in the 1^st^ week whereas late post-traumatic seizures or PTE started anywhere from months 1 to 3 after injury. We performed video-EEG monitoring for 1 week at a time in the first week, and then again at months 1, 2 and 3 and it is possible that more animals had seizures which we simply missed during times when EEG was not recorded. It is important to note that only 1 animal out of 15 had acute as well as chronic post-traumatic seizures, though this may be an under estimate as some animals had chronic recordings only. Others have reported “non-convulsive seizures” as well as “absence-like” events in Sprague-Dawley rats^24^ that had CCI. We did not find any electro-clinical seizures in our sham controls or NC controls but we saw isolated spikes, spike runs or absence-like spike-wave discharges in the sham injury animals, similar to that of previous reports^24, 28^. Previously, absence-like spike wave discharges were reported in sham group of Sprague-Dawley^28^ and in outbred Sprague-Dawley and Long-Evans rats or even wild-caught rats^29^. Given that manual detection of interictal spike counts can be very labor intensive, we used an automated method of analysis according to methods reported by our group previously in a repeated low-dose Kainic acid model^21^ and automated analysis of spike-wave discharges in a model of absence epilepsy ^22^. Interestingly, we found high amplitude events in all groups including NC controls, similar to what we observed in saline treated C57BL6 mice in the Kainate model^21^. However, high amplitude events with a unique morphology that was highly specific to the CCI group were also found in the sham but not in the NC group. In the Kainate model, animals with electro-clinical seizures had a much higher index of high amplitude events suggesting that they are more likely to occur in animals that eventually develop seizures. In our sample, even though the sham group had TBI-related events similar to those found in the CCI group, we did not observe any electro-clinical seizures. In addition, CCI animals that developed seizures did not have any greater frequency of the high amplitude events compared to those that did not develop acute or chronic post-traumatic seizures.

Many different studies reported sleep-wake disturbances in rodent models of TBI including the CCI model^30–35^, lateral or midline fluid percussion injury model^16, 36–40^ and in the weight drop model^41, 42^, but the data are conflicting. Most of the studies in the CCI model were done acutely, within the first week or two. Changes in sleep architecture reported in CCI models include increased NREM sleep, shorter wake bouts, increased sleep-wake fragmentation ^30, 31^; increased latency to reach peak sleep^32^ and hypersomnia acutely^33^. One study reported no changes in sleep-wake patterns after TBI^39^ and another reported increased mean percent time spent sleeping in the first week but not 2-5 weeks post injury^39^. Thus, in the TBI literature, no consistent pattern of sleep-wake disturbances have been seen and there is a dearth of studies that systematically examined these changes longitudinally. We, on the other hand, did perform longitudinal analyses but did not find any sleep-wake disturbances that are attributable to solely to the CCI itself at acute or chronic time points. We however found the normal expected oscillation of sleep/wake where there was less time awake and more in sleep during lights-on and the opposite during lights-off suggesting that the normal circadian distribution of sleep was preserved in all groups. One exception we found was that both sham and CCI groups spent greater time in NREM in the 4^th^ bin (6:30 pm to 10:30 pm, which is the first 4 hours of rodent wake time) suggesting that craniotomy itself resulted in “early morning sleepiness”. Furthermore, shorter wake or NREM bouts were seen in animals that had CCI or sham injury both acutely (week 1 after injury) or chronically (1-2 months after injury), again suggesting that any “sleep fragmentation” was seen in all animals that had a craniotomy rather than CCI alone.

Sleep spindle characteristics have been described as a potential biomarker of PTE in previous studies but data are conflicting^43, 44^. One study reported increased spindle frequency and duration in mice that had TBI compared to their sham controls^43^. However, another group reported shorter duration of spindles during transition from NREM to REM sleep in rats with TBI compared to a sham group and even among TBI, those that developed seizures had shorter duration of spindles compared to those that had no post-traumatic seizures^43^. The same group later reported that spindle characteristics could be potential predictors of PTE^45^. We also demonstrate that the normal 24-hour oscillation of spindles across lights-on and lights-off^46^ is preserved in the CCI model here, where they progressively increase in number during lights-on and decline during lights-off (as opposed to NREM delta power) regardless of whether they had a craniotomy or not. The only finding we had with respect to spindle characteristics was a relative decrease in spindle duration and power in sham and CCI groups compared to NC controls both acutely and chronically. We did not find any single spindle characteristic that was predictive of PTE.

One study examined NREM delta power sequentially at intervals, from acute time point to 30 days after injury in a CCI model and reported that NREM delta power increases transiently after CCI^35^. The NREM delta power is a marker of homeostatic sleep pressure and in humans it normally oscillates across the 24 hours, where it declines with sleep (as the homeostatic sleep drive decreases) and increases during the wake period as the homeostatic drive for sleep (or sleep pressure) increases^47^. It was previously demonstrated that in rodents including CD-1 mice, NREM delta power decreases progressively during lights-on when rodents sleep and increases during lights off as they spend more time awake^45, 48–49^. In our model, interestingly, we did not find these dynamics or the oscillation in CCI or sham or NC controls acutely or chronically. More importantly, however, we found several changes in mean NREM delta power. First is that there is an increase in the mean NREM delta power that followed injury severity from NC to sham to CCI especially during the first week. Chronically, the mean delta power in NREM sleep remained significantly higher in sham and CCI groups than NC control. Lastly, when we parsed CCI group into those with and without seizures, we noted a difference where CCI animals with seizures had mean NREM delta power that is similar to levels seen in NC controls at a chronic time point. The presence of seizures or interictal spikes themselves can result in increased synaptic strengthening that can contribute to increased NREM delta power^50^. Thus, in the CCI model, the presence of epileptiform activity seen in both sham and CCI groups might have possibly led to the increased NREM delta power in both. Given that NC animals had far fewer high amplitude interictal events, it is not surprising that NREM delta power is significantly lower in NC than the other groups. While seizures, especially those that are generalized and convulsive, can also increase NREM delta power, we saw no difference between those that did or did not develop seizures. This is contrary to conventional wisdom, but the reason behind reduced NREM delta power at the chronic time point in those that developed post-traumatic seizures is unclear but it is possible that interictal spikes rather than seizures have a far greater impact on increasing NREM delta power and should be pursued in future studies^50^.

Our data show that sham injury animals also had interictal phenomena as well as some sleep-wake changes similar to what was seen in the CCI group. During the study, because we saw the interictal phenomena in the sham group, we decided to employ an NC control group but also followed with an imaging study in sham and CCI cohorts, to determine if sham group had any extent of injury. Our CT imaging data showed that indeed animals with sham injury also had a visually evident lesion to the extent of the craniotomy though on quantification, the lesion was substantially smaller than in the CCI group (Figure S3). Taken together with the findings on sleep-wake changes and interictal phenomena in both CCI and sham groups, one speculation is that perhaps that the craniotomy itself in sham animals resulted in some degree of cerebral injury that contributed to these findings. The reasons behind why interictal epileptiform phenomena occurred in sham animals in previous reports is not clear and it is unknown whether any imaging was performed in this group to determine if they had any degree of cerebral injury.

There are several limitations to our interpretation of the data. First, our sample sizes are small. We performed EEG recording for a week at a time acutely after the injury, and again at months 1, 2 or 3. We may have missed seizures or even non-convulsive events when animals were not being recorded. We have not quantified all the different types of epileptiform activity using automated algorithms. Sleep scoring in 4-second epochs is very labor intensive and we only analyzed sleep data in a proportion of animals at week 1 and month 1 or 2. A larger sample size is probably more desirable. Using automated methods of sleep analysis may be a potential strategy in the future to process large amounts of data. Future studies could also determine the chronological changes in sleep or epileptiform activity after TBI. All the sleep analyses were performed in groups of animals and future experiments may be designed to sequentially perform studies in each mouse before and after CCI, such that every animal becomes its own control allowing comparing not only group differences but also within-animal differences. It is likely that our surgical technique resulted in some degree of cerebral injury in the sham group of animals, which would be a weakness of the study. A histopathology study might have yielded definitive information of any degree of brain injury in the sham group, which was not performed here and therefore a weakness. In the future, perhaps employing a no craniotomy control only may be more prudent.

## Conclusions

In conclusion, our data suggest that CCI results in acute post-traumatic seizures or PTE in a proportion of animals. While no seizures were seen in sham or NC groups, interictal epileptiform activity in the form of isolated interictal spikes, spike runs or “absence-like” spike-wave discharges were seen in both CCI and sham injury groups. Furthermore, high amplitude interictal events were seen in all groups, though the ones with high specificity for TBI were seen to a far greater extent in those with CCI compared to NC. In addition, the main sleep disruptions noted include shorter bout length for wake and NREM, reduced spindle duration or power as well as increased NREM delta power in sham and CCI groups, but no single sleep architecture characteristic in our sample was predictive of PTE. Given that imaging also showed a small volume lesion in sham animals to the extent of the craniotomy, we speculate that sham animals suffered a milder form of brain injury during the craniotomy itself that contributed to our findings. A pathological examination could resolve this more definitively and should be consideration for future studies. The significance of decline in NREM delta power in animals that developed PTE at the chronic time point is unclear but further work is needed to understand how NREM delta power evolves after TBI and its relation to development of PTE. Given the changes we observed in interictal phenomena, sleep-wake patterns, spindles and NREM delta power in Sham injury animals, using animals with no craniotomy only as controls is a strong consideration for future studies. These studies are relevant to further research in TBI models, to develop a sleep biomarker for PTE. The work is also relevant to humans with TBI as monitoring sleep phenotypes may predict risk, but may also help develop therapies to prevent post-traumatic epilepsy.

## Supporting information

Suppementary Figures and Tables

## Acknowledgement

Authors would like to thank Dr. Richard McNally’s Lab for providing the Matlab scripts for analysis sleep spindles.

## Financial or non-financial disclosures

None

