## Supplementary material for "Sleep-wake Characteristics in a Mouse Model of Severe Traumatic Brain Injury: Relation to Post-Traumatic Epilepsy": Suppementary Figures and Tables

Figure S1: Sleep scoring in 4 second epochs using Sirenia sleep software is shown. Each panel has a frontal EEG (first tracing), a parietal EEG (second tracing) and an EMG electrode (third tracing). EEG is band pass filtered from 1-70 Hz and EMG was band pass filtered between 5-500Hz prior to scoring sleep. Panel A shows a wake pattern with low amplitude EEG and increased EMG tone as well as some movement. Panel B shows a NREM sleep pattern with high amplitude frontal EEG and low EMG tone. Panel C shows low amplitude EEG and low EMG tone.

Figure S2: Data of sleep bout analysis is shown for Wake (A and D), NREM (B and E) and REM (C and F) in week 1 and month 1 or 2. Mean bout length and 95%CI for each group as well as p value is shown in each panel.

Figure S3: Data for number of bouts (episodes) of each vigilance state is shown among NC control, Sham injury or TBI group (Mean±SEM is shown). Significance in group differences was indicated by an *.

Figure S4: Ex vivo CT imaging data is shown for **(A, top row)** five brains from animals that had TBI. Notice the lesion produced by TBI on the right hemisphere (left side of image). **(B, bottom row)** five brains from Sham animals that received a craniotomy without TBI. **(C)** differences in volume of lesion at the TBI/craniotomy site is statistically significant. (Sham: 0.82 ± 0.68; TBI: 10.49 ± 7.2 $mm^3$; p = 0.039).

Table S1: Data is shown for time spent awake only across the 24 hours in six 4-hour bins among NC control, Sham and TBI at week 1 and Month 1 or 2. Analysis using repeated measures of ANOVA showed main effect of time and an oscillation pattern both in week 1 [F(5, 17)=6.74; p=0.003] and at month 1 or 2 [F(5, 17)=6.78; p=0.003], where the time spent in Wake was lower in Bins 1 and 2 (lights on) and gradually increased in Bins 3 and 4) during lights off before declining in the last bin (prior to lights on).

Table S2: Results from post-hoc multiple comparisons tests associated with the ANOVA analyses of NREM nDelta power graphically represented in Figure 5 are shown here. All values are mean ± 95% confidence interval. Panels A-C summarize the binned analysis for NC control, SHAM and TBI groups at acute and chronic timepoints, showing individual means (panel A), bin values of all groups (panel B), and group means (panel C). Panels D-F summarized the binned analysis of TBI animals, with or without post-traumatic seizures, at acute and chronic timepoints, showing individual means (panel D), mean bin values of both groups (panel E), and group means (panel F).

<u>Figure S1:</u>

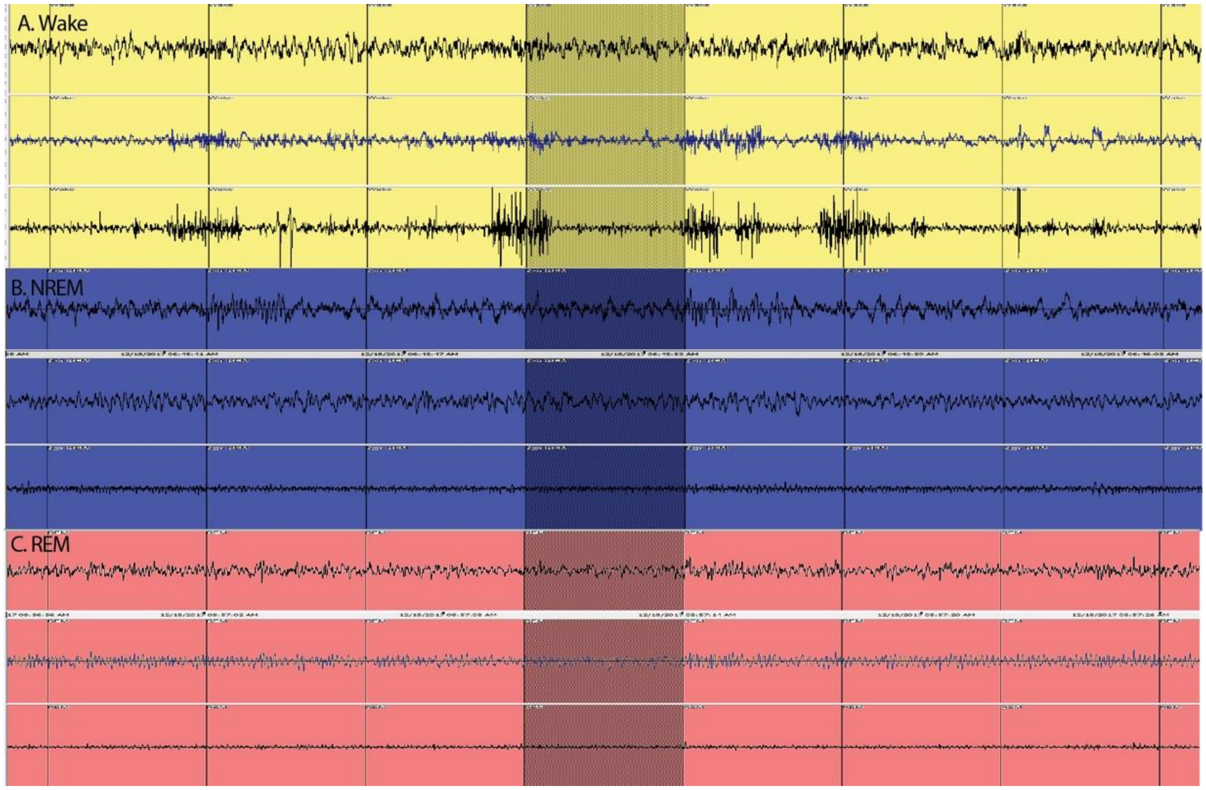

Figure S2:

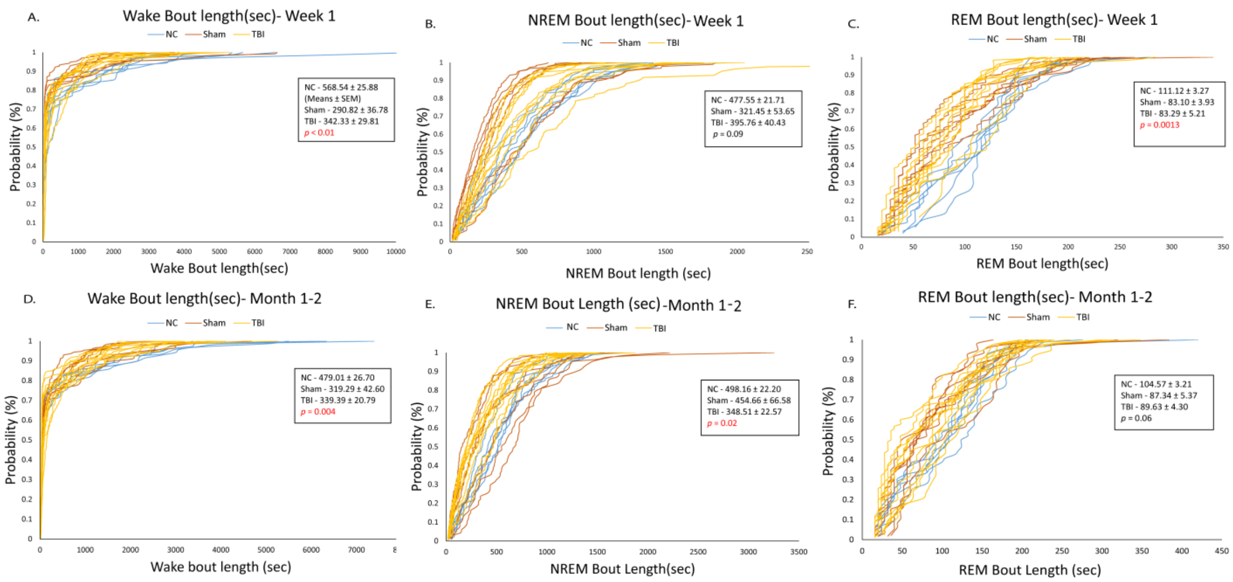
A.
Wake Bout length(sec)- Week 1
NC Sham TBI
NC - 568.54 ± 25.88
(Means ± SEM)
Sham - 290.82 ± 36.78
TBI - 342.33 ± 29.81
p < 0.01
Probability (%)
Wake Bout length(sec)
B.
NREM Bout length(sec)- Week 1
NC Sham TBI
NC - 477.55 ± 21.71
Sham - 321.45 ± 53.65
TBI - 395.76 ± 40.43
p = 0.09
Probability (%)
NREM Bout length (sec)
C.
REM Bout length(sec)- Week 1
NC Sham TBI
NC - 111.12 ± 3.27
Sham - 83.10 ± 3.93
TBI - 83.29 ± 5.21
p = 0.0013
Probability (%)
REM Bout length(sec)
D.
Wake Bout length(sec)- Month 1-2
NC Sham TBI
NC - 479.01 ± 26.70
Sham - 319.29 ± 42.60
TBI - 339.39 ± 20.79
p = 0.004
Probability (%)
Wake bout length (sec)
E.
NREM Bout Length (sec) -Month 1-2
NC Sham TBI
NC - 498.16 ± 22.20
Sham - 454.66 ± 66.58
TBI - 348.51 ± 22.57
p = 0.02
Probability (%)
NREM Bout Length(sec)
F.
REM Bout length(sec)- Month 1-2
NC Sham TBI
NC - 104.57 ± 3.21
Sham - 87.34 ± 5.37
TBI - 89.63 ± 4.30
p = 0.06
Probability (%)
REM Bout Length (sec)

Figure S3:

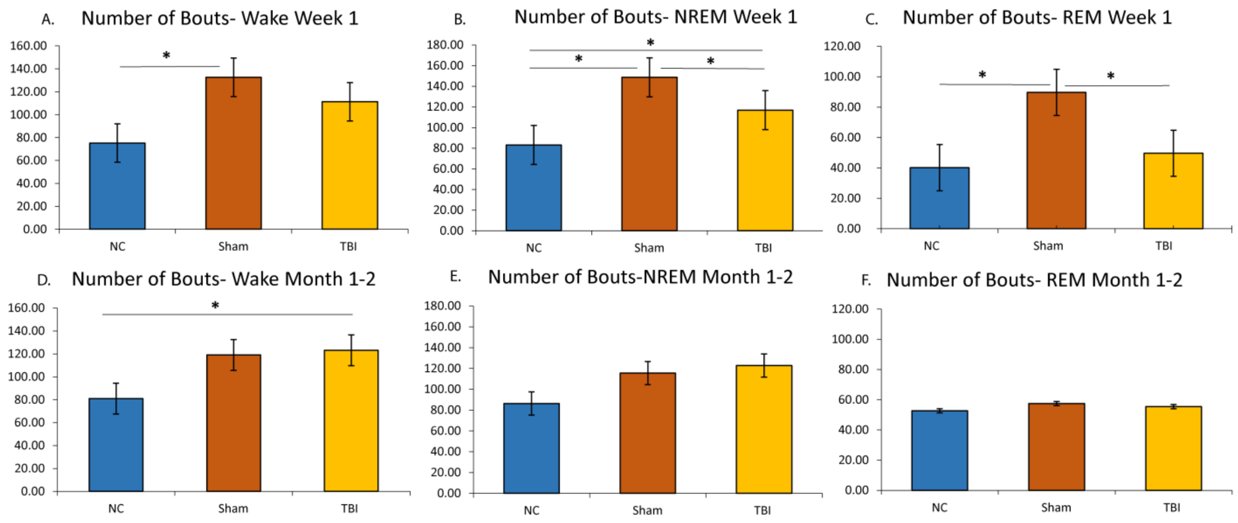
A.
Number of Bouts- Wake Week 1
160.00
140.00
120.00
100.00
80.00
60.00
40.00
20.00
0.00
*
NC
Sham
TBI
B.
Number of Bouts- NREM Week 1
180.00
160.00
140.00
120.00
100.00
80.00
60.00
40.00
20.00
0.00
*
*
*
NC
Sham
TBI
C.
Number of Bouts- REM Week 1
120.00
100.00
80.00
60.00
40.00
20.00
0.00
*
*
NC
Sham
TBI
D.
Number of Bouts- Wake Month 1-2
160.00
140.00
120.00
100.00
80.00
60.00
40.00
20.00
0.00
*
NC
Sham
TBI
E.
Number of Bouts-NREM Month 1-2
180.00
160.00
140.00
120.00
100.00
80.00
60.00
40.00
20.00
0.00
NC
Sham
TBI
F.
Number of Bouts- REM Month 1-2
120.00
100.00
80.00
60.00
40.00
20.00
0.00
NC
Sham
TBI

Figure S4:

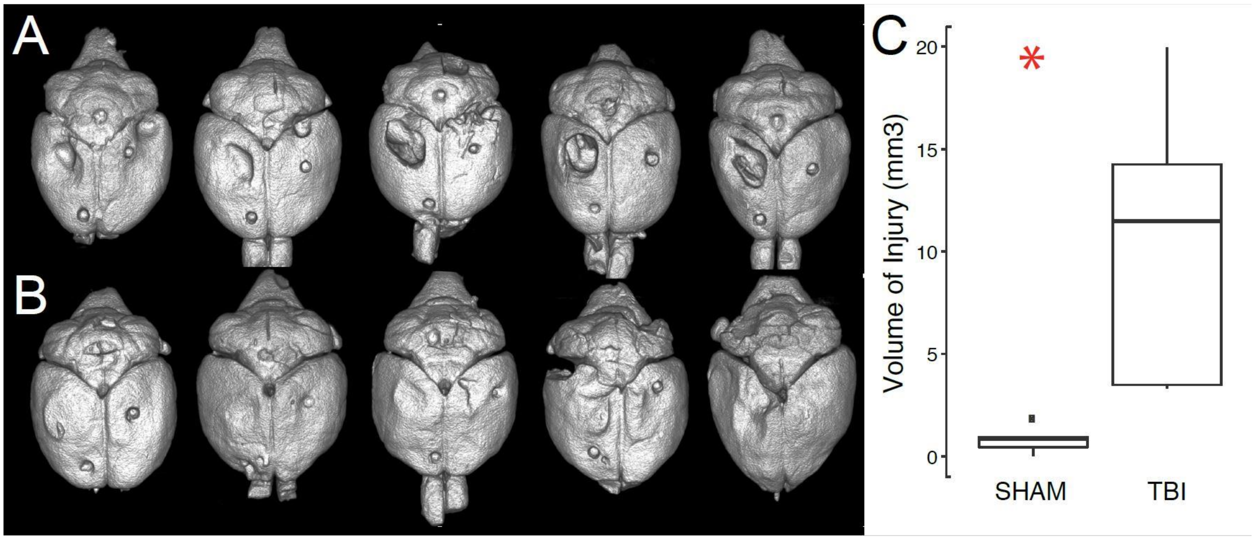

| Table S1: Mixed-Model ANOVA statistics of time spent in each Vigilance state in minutes, in each of the 4-hour bins | | | |
|---|---|---|---|
| Week 1 of Recording | | | |
| Time Bin | NC control | Sham | TBI |
| Bin 1 | 90.46±5.43 | 89.83±19.06 | 89.25±33.99 |
| Bin 2 | 79.31±3.65 | 77.99±18.22 | 82.88±21.46 |
| Bin 3 | 117.09±6.0 | 111.24±26.7 | 112.5±34.43 |
| Bin 4 | 131.89±5.21 | 124.74±24.2 | 120.02±14.54 |
| Bin 5 | 116.11±7.07 | 105.91±29.6 | 110.37±36.15 |
| Bin 6 | 96.60±7.04 | 79.33±28.73 | 99.16±38.53 |
| Month 1 or 2 of recording | | | |
| Bin 1 | 85.82±8.36 | 73.43±21.06 | 93.18±17.18 |
| Bin 2 | 62.37±19.12 | 67.12±19.72 | 72.77±18.14 |
| Bin 3 | 104.42±13.76 | 104.44±13.76 | 122.46±18.53 |
| Bin 4 | 139.53±10.87 | 117.90±18.21 | 152.68±18.54 |
| Bin 5 | 131.45±18.60 | 103.94±16.80 | 117.55±18.08 |
| Bin 6 | 115.83±27.41 | 98.68±20.82 | 119.15±19.09 |

Table S2:

A

| | Bin | Acute | Chronic |
|---|---|---|---|
| NC | 1 | 0.9331 ± 0.0079 | 0.8753 ± 0.0107 |
| | 2 | 1.0051 ± 0.0076 | 0.9086 ± 0.0100 |
| | 3 | 0.9736 ± 0.0091 | 0.8627 ± 0.0113 |
| | 4 | 0.9305 ± 0.0109 | 0.7645 ± 0.0129 |
| | 5 | 0.8779 ± 0.0093 | 0.7327 ± 0.0124 |
| | 6 | 0.8613 ± 0.0084 | 0.6904 ± 0.0116 |
| SHAM | 1 | 1.0709 ± 0.0086 | 1.0563 ± 0.0105 |
| | 2 | 1.0721 ± 0.0083 | 1.1182 ± 0.0106 |
| | 3 | 1.1122 ± 0.0096 | 1.0492 ± 0.0116 |
| | 4 | 1.1242 ± 0.0102 | 1.0227 ± 0.0122 |
| | 5 | 1.0557 ± 0.0088 | 0.9152 ± 0.0115 |
| | 6 | 1.0291 ± 0.0079 | 0.9313 ± 0.0112 |
| TBI | 1 | 1.3533 ± 0.0079 | 1.1131 ± 0.0099 |
| | 2 | 1.3320 ± 0.0075 | 1.1236 ± 0.0094 |
| | 3 | 1.3501 ± 0.0082 | 1.0624 ± 0.0109 |
| | 4 | 1.3261 ± 0.0086 | 1.0695 ± 0.0116 |
| | 5 | 1.2731 ± 0.0081 | 1.0108 ± 0.0103 |
| | 6 | 1.2374 ± 0.0078 | 0.9475 ± 0.0101 |

B

| Bin | Acute | Chronic |
|---|---|---|
| 1 | 1.1191 ± 0.0047 | 1.0149 ± 0.0060 |
| 2 | 1.1364 ± 0.0045 | 1.0501 ± 0.0058 |
| 3 | 1.1453 ± 0.0052 | 0.9914 ± 0.0065 |
| 4 | 1.1269 ± 0.0057 | 0.9522 ± 0.0071 |
| 5 | 1.0689 ± 0.0050 | 0.8863 ± 0.0066 |
| 6 | 1.0426 ± 0.0046 | 0.8564 ± 0.0063 |

C

| Group | Acute | Chronic |
|---|---|---|
| NC | 0.9303 ± 0.0036 | 0.8057 ± 0.0047 |
| SHAM | 1.0774 ± 0.0036 | 1.0155 ± 0.0046 |
| TBI | 1.3120 ± 0.0033 | 1.0545 ± 0.0042 |

D

| | Bin | Acute | Chronic |
|---|---|---|---|
| Seizures + | 1 | 1.3387 ± 0.0133 | 1.0120 ± 0.0146 |
| | 2 | 1.3663 ± 0.0131 | 1.0599 ± 0.0139 |
| | 3 | 1.3262 ± 0.0153 | 0.9559 ± 0.0156 |
| | 4 | 1.3237 ± 0.0162 | 0.9623 ± 0.0177 |
| | 5 | 1.2663 ± 0.0151 | 0.8996 ± 0.0150 |
| | 6 | 1.2126 ± 0.0135 | 0.8415 ± 0.0148 |
| Seizures - | 1 | 1.3632 ± 0.0110 | 1.2134 ± 0.0146 |
| | 2 | 1.3115 ± 0.0102 | 1.1848 ± 0.0137 |
| | 3 | 1.3622 ± 0.0109 | 1.1781 ± 0.0163 |
| | 4 | 1.3273 ± 0.0113 | 1.1627 ± 0.0165 |
| | 5 | 1.2764 ± 0.0106 | 1.1243 ± 0.0151 |
| | 6 | 1.2527 ± 0.0106 | 1.0556 ± 0.0149 |

E

| Bin | Acute | Chronic |
|---|---|---|
| 1 | 1.3510 ± 0.0086 | 1.1127 ± 0.0103 |
| 2 | 1.3389 ± 0.0083 | 1.1224 ± 0.0098 |
| 3 | 1.3442 ± 0.0094 | 1.0670 ± 0.0113 |
| 4 | 1.3255 ± 0.0099 | 1.0625 ± 0.0121 |
| 5 | 1.2714 ± 0.0093 | 1.0119 ± 0.0106 |
| 6 | 1.2326 ± 0.0086 | 0.9485 ± 0.0105 |

F

| Group | Acute | Chronic |
|---|---|---|
| Seizure + | 1.3056 ± 0.0059 | 0.9552 ± 0.0063 |
| Seizure - | 1.3156 ± 0.0044 | 1.1531 ± 0.0062 |
